# Paralog editing tunes rice stomatal density to maintain photosynthesis and improve drought tolerance

**DOI:** 10.1101/2021.12.21.473329

**Authors:** Nicholas G. Karavolias, Dhruv Patel-Tupper, Kyungyong Seong, Michelle Tjahjadi, Gloria-Alexandra Gueorguieva, Jaclyn Tanaka, Ana Gallegos Cruz, Samantha Liebermann, Lillian Litvak, Douglas Dahlbeck, Myeong-Je Cho, Krishna K. Niyogi, Brian J. Staskawicz

**Affiliations:** University of California, Berkeley; Innovative Genomics Institute; University of California, Berkeley; Innovative Genomics Institute; UC Berkeley, Howard Hughes Medical Institute; Lawrence Berkeley National Laboratory, Molecular Biophysics and Integrated Bioimaging Division

## Abstract

Rice (*Oryza sativa*) is of paramount importance for global nutrition, supplying at least 20% of global calories. However, water scarcity and increased drought severity are anticipated to reduce rice yields globally. Rice stomatal developmental genetics were explored as a mechanism to improve drought resilience while maintaining yield under climate stress. CRISPR/Cas9-mediated knockouts of EPFL10 and STOMAGEN yielded lines with c. 80% and 25% of wild-type stomatal density, respectively. *epfl10* lines with moderate reductions in stomatal densities are able to conserve water to similar extents as *stomagen* lines, but do not suffer from the concomitant reductions in stomatal conductance, carbon assimilation, or thermoregulation observed in *stomagen* knockouts. Moderate reductions in stomatal densities achieved by editing EPFL10 may be a climate-adaptive approach in rice that can safeguard yield. Editing the paralog of STOMAGEN in other species may provide a means to tune stomatal density in agriculturally important crops beyond rice.

## Introduction

Prolonged periods of drought and increased desertification are anticipated to become more prevalent in the next century ^1^. Climate change modeling predicts increases in global temperatures by 2-4 °C by the end of the 21^st^ century. Increased temperatures alone and in combination with limited water will negatively impact crop yields ^2–4^. The development of climate change adapted crops is essential to maintain crop yields in the face of rapid global population growth and worsening climates.

Rice (*Oryza sativa*) is one of the most widely consumed crops globally and trails only wheat and maize in area harvested ^5^. Originally domesticated in semi-aquatic habitats, rice is especially sensitive to drought relative to other C3 cereal crops ^6–8^. Future water limitations may necessitate transitions of fully flooded paddy fields to water-saving production schemes ^4^. Rain-fed production, which comprises about 45% of total rice grown, is particularly susceptible to drought as a result of unpredictable precipitation ^9–11^. Furthermore, most regions where irrigated rice is produced are currently experiencing or projected to experience water scarcity ^10^. Thus, all rice, regardless of production methods, would benefit from improvements that maintain yields with less water.

Stomata are at the nexus of plants and the atmosphere. They facilitate gaseous exchanges of carbon dioxide, oxygen, and water vapor. Most water loss in crops occurs via transpiration from the stomata ^12^. Breeding efforts have shown drought adaptation in rice was facilitated in part by reductions of stomatal density ^13^. Thus, opportunity exists, especially in non-adapted cultivars, to fine tune stomatal density reductions.

The developmental biology of stomata has been studied extensively in myriad species^14–17^. Despite similar sequence identities, EPIDERMAL PATTERNING FACTORs (EPFs) and EPF-LIKE proteins (EPFLs) mediate opposing downstream stomatal development responses. EPF1 and EPF2 function as negative regulators expressed in stomatal-lineage cells. In contrast, EPFL9, also known as STOMAGEN, is a positive regulator of stomatal development that is dynamically expressed in the mesophyll ^18–24^. These mobile peptides regulate cell fate transitions and cell divisions to ensure proper spacing and number of stomata ^18,21–25^.

EPFL9 is composed of three distinct regions: an N-terminal signal peptide region, a pro-peptide region, and a C-terminal cysteine-rich active peptide region ^18,26^. The full-length peptide is processed in vivo to yield a 45-residue C-terminal active peptide ^18,25^. The active peptide encoded by EPFL9 possesses the conserved cysteine residues of EPF1 and EPF2 and binds the same ERECTA (ER)-family receptors and co-receptor TOO MANY MOUTHS (TMM) in *Arabidopsis thaliana* ^18,21,25,27^. A knockout of EPFL9 in rice using CRISPR/Cas9 and CRISPR/Cpf1 yielded an eight-fold reduction in abaxial stomatal density in the IR64 background ^19^.

Interestingly, EPFL9 appears to have undergone a duplication event in grasses ^27^. Transgenic overexpression of rice STOMAGEN and its duplicate, previously named EPFL9-2, in *A. thaliana* revealed a shared, though attenuated, function of EPFL9-2 as a positive regulator of stomatal development when ectopically expressed ^28^. Ectopic expression of *Brachypodium distachyon* and *Triticum aestivum* (wheat) STOMAGEN and STOMAGEN paralogs in *A. thaliana* resulted in similar stomatal density increases ^29^. In contrast, overexpression of negative regulators of stomatal development reduced stomatal density. Stomatal density reductions improved water use efficiency in *A. thaliana*, wheat, barley, and rice ^30–35^. However, all stomatal density reductions achieved by overexpressing negative regulators of stomatal development to any extent also reduced stomatal conductance and carbon assimilation under physiologically relevant light conditions ^30,31,33^. For example, rice lines overexpressing EPF1 to reduce stomatal densities exhibited lower stomatal conductance and carbon assimilation at all light conditions that exceeded 1,000 μmol photons m^−2^ s^−1 33^. These reductions may be detrimental as stomatal conductance has been shown to be essential for crop productivity ^36–43^. In rice specifically, higher stomatal conductance has been associated with greater rates of leaf photosynthesis ^37,39,44^. Also of note, all previously tested strategies relied on the use of constitutive transgenic overexpression, lacking the tissue specificity and gene dosage that may regulate stomatal density phenotypes *in vivo*.

Gene editing of rice STOMAGEN and its duplicate were used as an alternative, non-transgenic, approach to tune stomatal density. Gene duplication events can provide genetic material for functional novelty ^47^, but may also disrupt optimal levels of gene expression, requiring significant evolutionary time to stabilize ^48^. Gene editing of paralogs can provide a potent and straightforward mechanism to generate desirable variation in a wide array of traits. For example, knockout alleles of individual genes in a set of paralogs underlying disease resistance and yield in tomato and maize, respectively, has demonstrated the utility of paralog gene editing for generating desirable trait variation ^49,50^.

Here, we report novel characterization of the rice STOMAGEN paralog EPFL9-2, subsequently referred to as EPFL10, and its relationship to stomatal development. Furthermore, we explore the effect of the reductions in stomatal density in *stomagen* and *epfl10* mutants on stomatal conductance, carbon assimilation, thermal regulation, water conservation, and yield in varying water regimes. We describe the use of paralog editing for achieving desirable variation in traits of interest and the implications of this work on future gene-editing strategies for improved water-use efficiency.

## Methods

### Plant Growth conditions

Rice cultivar Nipponbare (*O. sativa* ssp. japonica) seeds were germinated and grown for 8 days in a petri dish with 20 mL of water in a Conviron growth chamber at 28°C for day-length periods of 16 hours in 100 μmol photons m^−2^ s^−1^ of light and 80% relative humidity. Seedlings were transferred to a soil mixture comprised of equal parts turface (https://www.turface.com/products/infield-conditioners/mvp) and sunshine mix #4 (http://www.sungro.com/professional-products/fafard/).

Germinated seedlings used for stomatal phenotyping and growth chamber physiological assays were transferred to 10 cm, 0.75 L McConkey tech square pots and placed in growth chambers at 28°C for day-length periods of 16 hours in 400 μmol photons m^−2^ s^−1^ of light and 80% relative humidity.

Plants designated for yield trials, greenhouse physiological assays, and stomatal aperture measurements were moved to the greenhouse with temperature setpoints of 27°C day/22°C night at ambient light conditions in February 2020 with day lengths of 12 hours in 15.2 cm, 1.8 L pots. All plants were fertilized with 125 mL of 1% w/v iron solution one-week post-transplant. 1 L of 5% w/v JR Peter’s Blue 20-20-20 fertilizer (https://www.jrpeters.com/) was added to each flat at 3- and 11-weeks post-germination.

### Yield and water regimes

Adapted from Caine et. al ^33^, grain and biomass yield in three watering regimes were tested: well-watered, vegetative drought, and reproductive drought. Well-watered flats were kept flooded for the entirety of the growth period. Vegetative drought was imposed by removing all water from flats containing pots for 7 days starting on day 28 after germination and for 9 days at day 56 after germination. In reproductive drought, water was removed from flats for 4 days at day 98 when panicles were undergoing grain filling. All grain and aboveground biomass from well-watered, vegetative and reproductive drought plants were harvested after 167 days, 177 days, and 181 days, respectively. Biomass measurements were completed on samples dried at 60°C for three days prior to weighing.

### Generation of edited lines

Guides for targeting EPFL10 and STOMAGEN were selected to minimize off-target effects and maximize on-target efficiency in the first exon of the coding region. Guide sequences were selected using CRISPR-P 2.0 (http://crispr.hzau.edu.cn/CRISPR2/). Forward and reverse strand guide sequences with relevant sticky ends amenable for Golden Gate cloning were ordered from Integrated DNA Technology (IDTdna.com). Equal volumes of 10 mM primers were annealed at room temperature. Golden Gate cloning was used to insert guides into the PeGM entry clone containing the tracrRNA and U3 promoter. LR clonase reactions were used to insert the entry clone into destination vectors for biolistic transformation and *Agrobacterium*-mediated transformation. *epfl10* lines were produced via *Agrobacterium*-mediated transformation and *stomagen* lines via biolistic transformation.

### Plant material and culture of explants

Mature seeds of rice (*O. sativa* L. japonica cv. Nipponbare) were de-hulled, and surface-sterilized for 20 min in 20% (v/v) commercial bleach (5.25% sodium hypochlorite) and 1% of Tween 20. Three washes in sterile water were used to remove residual bleach from seeds. De-hulled seeds were placed on callus induction medium (CIM) medium [N6 salts and vitamins ^51^, 30 g/L maltose, 0.1 g/L myo-inositol, 0.3 g/L casein enzymatic hydrolysate, 0.5 g/L L-proline, 0.5 g/L L-glutamine, 2.5 mg/L 2,4-D, 0.2 mg/L BAP, 5 mM CuSO_4_, 3.5 g/L Phytagel, pH 5.8] and incubated in the dark at 28 °C to initiate callus induction. Six- to 8-week-old embryogenic calli were used as targets for transformation.

### *Agrobacterium*-mediated transformation

Embryogenic calli were dried for 30 min prior to incubation with an *Agrobacterium tumefaciens* EHA105 suspension (OD_600nm_ = 0.1) carrying the binary vector for editing rice EPFL10. After a 30 min incubation, the *Agrobacterium* suspension was removed. Calli were then placed on sterile filter paper, transferred to co-cultivation medium [N6 salts and vitamins, 30 g/L maltose, 10 g/L glucose, 0.1 g/L myo-inositol, 0.3 g/L casein enzymatic hydrolysate, 0.5 g/L L-proline, 0.5 g/L L-glutamine, 2 mg/L 2,4-D, 0.5 mg/L thiamine, 100 mM acetosyringone, 3.5 g/L Phytagel, pH 5.2] and incubated in the dark at 21°C for 3 days. After co-cultivation, calli were transferred to resting medium [N6 salts and vitamins, 30 g/L maltose, 0.1 g/L myo-inositol, 0.3 g/L casein enzymatic hydrolysate, 0.5 g/L L-proline, 0.5 g/L L-glutamine, 2 mg/L 2,4-D, 0.5 mg/L thiamine, 100 mg/L timentin, 3.5 g/L Phytagel, pH 5.8] and incubated in the dark at 28°C for 7 days. Calli were then transferred to selection medium [CIM plus 250 mg/L cefotaxime and 50 mg/L hygromycin B] and allowed to proliferate in the dark at 28°C for 14 days. Well-proliferating tissues were transferred to CIM containing 75 mg/l hygromycin B. The remaining tissues were subcultured at 3- to 4-week intervals on fresh selection medium. When a sufficient amount (about 1.5 cm in diameter) of the putatively transformed tissues was obtained, they were transferred to regeneration medium [MS salts and vitamins ^52^, 30 g/L sucrose, 30 g/L sorbitol, 0.5 mg/L NAA, 1 mg/L BAP, 150 mg/L cefotaxime] containing 40 mg/L hygromycin B and incubated at 26 °C, 16-h light, 90 μmol photons m^−2^ s^−1^. When regenerated plantlets reached at least 1 cm in height, they were transferred to rooting medium [MS salts and vitamins, 20 g/L sucrose, 1 g/L myo-inositol, 150 mg/L cefotaxime] containing 20 mg/L hygromycin B and incubated at 26 °C under conditions of 16-h light (150 μmol photons m^−2^ s^−1^) and 8-h dark until roots were established and leaves touched the Phytatray™ II lid (Sigma-Aldrich, St. Louis, MO, USA). Plantlets were then transferred to soil.

### Biolistic-mediated transformation

Embryogenic callus tissue pieces (3–4 mm) were transferred for osmotic pretreatment to CIM medium containing mannitol and sorbitol (0.2 M each). Four hours after treatment with osmoticum, tissues were bombarded as previously described ^53^ with modifications. Two mg of gold particles (0.6 μm), coated with 5 mg of plasmid DNA for editing rice STOMAGEN were divided equally among 10 macro-carriers and used for bombardment with a Bio-Rad PDS-1000 He biolistic device (Bio-Rad, Hercules, CA, USA) at 650 psi. Sixteen to 18 h after bombardment, tissues were placed on osmotic-free CIM and incubated at 28°C under dim light (10-30 μmol photons m^−2^ s^−1^, 16-h light). After 7 days, tissues were transferred to a selection medium (CIM containing 50 mg/l hygromycin B) and grown using the same procedure as described above, without timentin or cefotaxime supplemented in the media.

### Validation of edits

T_0_ plants targeted for edits in EPFL10 and STOMAGEN were evaluated using PCR to amplify the region of interest. PCR products were Sanger sequenced. Sequence data was analyzed using the Synthego ICE tool (https://ice.synthego.com/#/) to detect alleles present ^54^. Only lines with homozygous frame-shift mutations were retained for downstream experiments. T_2_ plants were used for experimental data collection to minimize somaclonal variation, which may have accumulated during tissue culture ^55,56^.

### Phenotyping stomatal density, size, and aperture

Stomatal densities were recorded from epidermal impressions of leaves using nail polish peels ^37^. Stomatal densities of eight biological replicates of each leaf were taken from the widest section of fully expanded leaves. Images were taken using a Leica DM5000 B epifluorescent microscope at 10x magnification. Three images were collected per stomatal impression and density per image was averaged. The number of stomata in a single stomatal band were counted and the area of each band was measured ^57^. Stomatal densities were calculated by dividing stomatal counts by stomal band area (mm^2^).

Epidermal peels of 21-day old plants were produced using a razor blade on the adaxial leaf to remove tissues above the abaxial epidermal layer. Images of individual stomata at 100x magnification were captured. Guard cell length was measured using ImageJ. 35 individual stomata from five biological replicates of each genotype were measured.

Stomatal aperture measurements were generated using epidermal peels of flag leaves from 85-day old plants. Leaves were harvested at 1:00 p.m. and peels were generated immediately. Epidermal peels were then fixed by submerging in 4% formaldehyde for 30 seconds using a method adapted by Eisele et al. ^58^. Images of 20 individual stomata from six biological replicates of each genotype were measured.

Confocal microscopy images were captured using a Zeiss LSM 710 on epidermal peels of each genotype stained with propidium iodide for 40 seconds and immediately washed in water. Images were processed using Bitplane’s Imaris.

### Quantifying STOMAGEN and EPFL10 transcript abundance

Total RNA was extracted from seedlings with the Qiagen Total RNAeasy Plant Kit at three developmental stages: eight days after germination, 15 days after germination from the basal 2.5cm above the leaf sheath of the fifth expanding leaf, and from the fully expanded leaf of 21-day-old leaves. RNA quality was validated on an agarose gel prior to reverse transcription using the QuantiTect™ reverse transcription kit to generate first-strand cDNA. Quantitative reverse transcription PCR was performed using FAST SYBR on Applied Biosystem’s QuantStudio 3 thermocycler. Relative expression levels were calculated by normalizing to the rice UBQ5 housekeeping gene (LOC_Os01g22490) ^59^. Primers used for qPCR listed in Supporting Table 3. Relative log fold expression was calculated using the 2 ^−ΔΔCT^ method using STOMAGEN in adult leaves as the control group.

### Determining methylation profile of genes of interest

Methylation profiles of rice genes of interest were viewed using the Plant Methylation Database (https://epigenome.genetics.uga.edu/PlantMethylome/) ^61^. Snapshots of CHH and CHG methylation 1.5 kb upstream of the start codon and 1.5 downstream of the stop codon were taken.

### Evolutionary analysis of STOMAGEN paralogs

Single-copy orthologs were searched using BUSCO v4.0.6 ^62^ and the viridiplantae_odb10 database for the species used in this study (Table S1). 82 orthologous groups present in at least 23 species were individually aligned with MAFFT v7.487 (--maxiterate 1000 --globalpair) ^63^. All multiple sequence alignments were concatenated, trimmed with TrimAl v1.4.rev15 (-gt 0.2)^64^ and then used to infer a species tree with FastTree v2.1.10 ^65^. The copy number variations of the STOMAGEN family was determined by searching for the stomagen domain (PF16851) from the protein annotation sets with hmmsearch v3.3 ^66,67^ or from the genomes with exonerate v2.2.0 ^68^ if genome annotations are absent. To understand the sequence variations of STOMAGEN and EPFL10 orthologs at the species and family level, we collected non-redundant *Oryza* or Poaceae species that have single copies of STOMAGEN or EPFL10 (Supporting Table S2). The stomagen domain of STOMAGEN or EPFL10 orthologs was aligned with MAFFT, and the filtered alignment was used to compute normalized Shannon’s entropy. Gaps were ignored.

### Photosynthesis and stomatal conductance assays

Light response curves were generated using a LI6800 infrared gas analyzer (LI-COR, Lincoln, NE, USA) with chamber conditions set to: leaf temperature 25°C; flow rate 500 μmol s^−1^; water vapor pressure deficit 1.8 kPA; and CO_2_ concentration of sample 400 μmol mol^−1^. Light intensity was first increased to 2000 μmol photons m^−2^s^−1^, and with steady-state waiting times of 5 to 10 minutes, subsequently decreased to 1500, 1200, 1000, 750, 500, 300, 200, 100, and 50 μmol photons m^−2^ s^−1^ light. Light was composed of at least 90% red light and at maximum 40 μmol photons m^−2^ s^−1^ blue light to match equipment specifications. Measurements were taken on fully expanded fifth leaves of 32-day-old plants grown in the greenhouse. Anatomical g_smax_ was calculated using the double end-corrected version of the Franks and Farquhar ^69^ equation from Dow et al. ^70^ as described in Caine et. al^33^.

Physiological assays in Figures 3a, 3c were conducted on fully expanded leaf 5 of 21-day old plants. Stomatal conductance and CO_2_ assimilation data for Fig 3c was captured using an infrared gas analyzer (LI6400XT, LI-COR, Lincoln, NE, USA) with chamber conditions set to: light intensity 1000 μmol photons m^−2^ s^−1^ (90% red light, 10% blue light); leaf temperature 27°C; flow rate 500 μmol s^−1^; relative humidity 40%; and CO_2_ concentration of sample 400 μmol mol^−1^.

### Thermal imaging

Thermal images were captured using a FLIR E8-XT Infrared Camera (FLIR-DIRECT, Wilmington, NC, USA). Images of well-watered and vegetative drought plants were taken 65 days after germination on the last day of the vegetative drought treatment. Images of reproductive drought plants were captured 102 days post-germination on the last day of the reproductive drought treatment. Images were captured between 1:00 p.m. and 2:00 p.m. to represent the effects of transpiration-mediated cooling during the hottest part of the day. Images were processed using FLIR Thermal Studio. Leaf temperatures of 4-6 leaves per biological replicate were quantified.

### Water loss

Two plants of identical genotype were placed in a 10“×10“ flat and were covered with aluminum foil to minimize evaporation from soil. Non-plant evaporation was estimated by measuring daily water loss from covered flats containing pots without plants. These control flats were distributed within the greenhouse assay area to reflect environmental variation. Daily water loss of each flat was calculated by taking the difference of flats with plants and the average of the flats without. Eight replicate flats for each genotype were measured daily from 70-77 days after germination.

### Graphs and statistics

All graphs were produced using the ggplot2 package in R studio ^71^. All statistics were calculated in R-studio using post-hoc tests in the PMCMRplus package for significance between groups ^72^.

## Results

### Evolution and regulation of STOMAGEN and EPFL10 suggests functional differences of paralogs

Previous work has suggested the STOMAGEN paralog EPFL10 (LOC_Os08g41360) may function as a weak, positive regulator of stomatal development ^28,29^, having arisen from a putative duplication event in the most recent common ancestor of the Poaceae species ^27^. We examined evolutionary divergence of STOMAGEN and EPFL10 that may explain such functional differences.The two paralogs have accumulated divergent point mutations. In rice, homology cannot be detected between STOMAGEN and EPFL10 across the N-terminal signal peptides and pro-peptide domain. The functional 45 C-terminal sequences show 73% sequence identity with 12 non-synonymous substitutions (Fig. 1C). These positions were mapped onto the existing complex structure of TMM, ERL1 and EPF1 (Fig. 1D), assuming the binding of STOMAGEN and EPFL10 may be similar to EPF1. Many positions highlighted with non-synonymous substitutions were in proximity to the receptors, possibly suggesting that the paralogs may have different binding affinity to the receptors.

**Figure 1.**
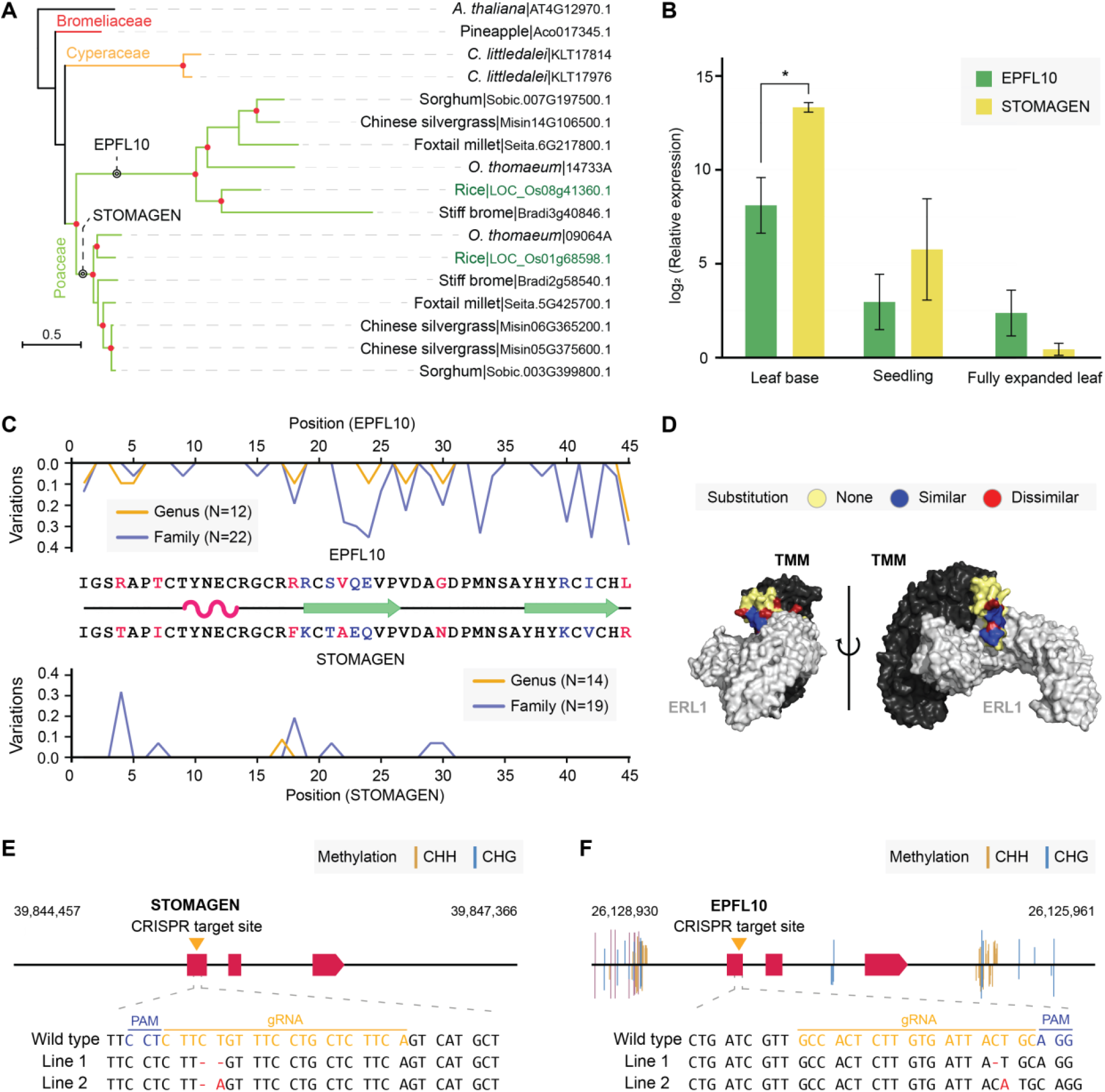
Evolution and regulation of STOMAGEN and EPFL10. **A**, Gene trees of the STOMAGEN family members in Poales. *Arabidopsis thaliana* was used as an outgroup. **B**, qRT-PCR determined expression levels of EPFL10 and STOMAGEN in fully expanded leaf, leaf base, and seedling. Bars represent means and error bars represent standard deviation from the mean. The asterisk indicates a significant difference between the means (P<0.05). **C**, Genus and family-level sequence variations of STOMAGEN and EPFL10 orthologs. The sequence variations represent normalized Shannon’s entropy, with 0 being no variations and 1 being complete variations. The secondary structure annotation originates from the experimentally determined structure of STOMAGEN (PDB: 2LIY) ^26^. The colored residues highlight variable positions between rice STOMAGEN and EPFL10. Substitutions to similar and dissimilar amino acids based on BLOSUM62 are indicated with blue and red. Alpha helix is represented by pink and beta sheets by green. **D**, The variable positions between STOMAGEN and EPFL10 mapped to the experimentally determined complex structure of EPF1, ERL1 (in gray) and TMM (in black) (PDB: 5XJO)^79^. Peptide sequence variations between EPFL10 and STOMAGEN with similar or dissimilar substitutions are indicated in blue and red, respectively. Complex is visualized from two orientations. **E** and **F**, The gene models of (E) STOMAGEN and (F) EPFL10 with the location of the CRISPR/Cas9 target sites indicated by the yellow triangle. Repressive CHH and CHG methylation marksare indicated for 1.5 kb up- and down-stream of the coding regions. The two unique homozygous edits generated in STOMAGEN and EPFL10 by CRISPR/Cas9 are shown in red with the PAM annotated in blue and guide sequence in yellow.

The phylogenetic tree also indicated that the two orthologous groups may have experienced differential selective pressure (Fig. 1A). To gain better insights, we calculated sequence variations of the STOMAGEN and EPFL10 orthologs with greater numbers of sequences collected from the *Oryza* species and the Poaceae family members (Fig. 1C; Table S1; Table S2). Consistent with the phylogeny, both species- and family-level sequence diversity was higher in EPFL10 orthologs, indicating relatively rapid divergence across the species. On the contrary, STOMAGEN orthologs have maintained a high level of sequence conservation across the species and family, suggesting their sequence evolution is strictly constrained. Furthermore, regulatory variations existed between STOMAGEN and EPFL10. Repressive CHG and CHH methylation marks vicinal to the EPFL10 coding region suggested regulatory restriction, while this was not observed for STOMAGEN. Concordantly, we observed that STOMAGEN mRNA abundance also greatly exceeded EPFL10 expression in leaf base tissues where stomatal development occurs and STOMAGEN and EPFL10 expression is greatest (Fig. 1B; Supporting Table S1). There is minimal expression of either transcript in adult leaf where stomatal complexes are fully matured (Fig 1B). Collectively, the sequence evolution and regulation of STOMAGEN and EPFL10 suggested possibly differential evolution and usage of the paralogs. This suggested that EPFL10 may be a better candidate to fine-tune stomatal development without abolishing the role of STOMAGEN, the primary positive regulator of stomatal development.

### Stomatal density and morphology are altered in knockout lines

To clarify the functional role of STOMAGEN and EFPL10 in regulating stomatal density *in vivo*, single and double mutant lines were generated in rice (*Oryza sativa* cv. Nipponbare). CRISPR/Cas9-mediated knockout of STOMAGEN and EPFL10 was achieved by targeting guides to the first exon of each gene to disrupt the open reading frame (Fig. 1E and 1F). Two unique homozygous knockout alleles were generated in EPFL10 or in STOMAGEN in the T0 generation using a single guide sequence adjacent to a PAM motif (Fig. 1E, 1F).

*epfl10* exhibited reductions in stomatal densities which represented 80% of wild-type densities, whereas *stomagen* possessed only 25% of wild-type densities in the fifth fully expanded adult leaf (Fig. 2A). Similar proportions of stomatal density reductions in mutants were measured in flag leaves and their adaxial leaf surfaces (Fig. 2B and 2C). Stomatal length was measured to determine if there was a relationship between stomatal density reductions and size increases in the Nipponbare background. The guard cell length of stomata in *stomagen* were longer relative to *epfl10* and wild type (Fig. 2D). Representative confocal microscopy captured stomatal density relationships in a single biological replicate of each genotype (Fig. 2E). The stomatal density of *stomagen epfl10* double mutants could not be distinguished from *stomagen* and were thus excluded from future physiological analyses (Fig. S1). Collectively, our data suggest that EPFL10 plays a functional role in rice stomatal formation, but the loss of STOMAGEN is epistatic to EPFL10-mediated stomatal development.

**Figure 2.**
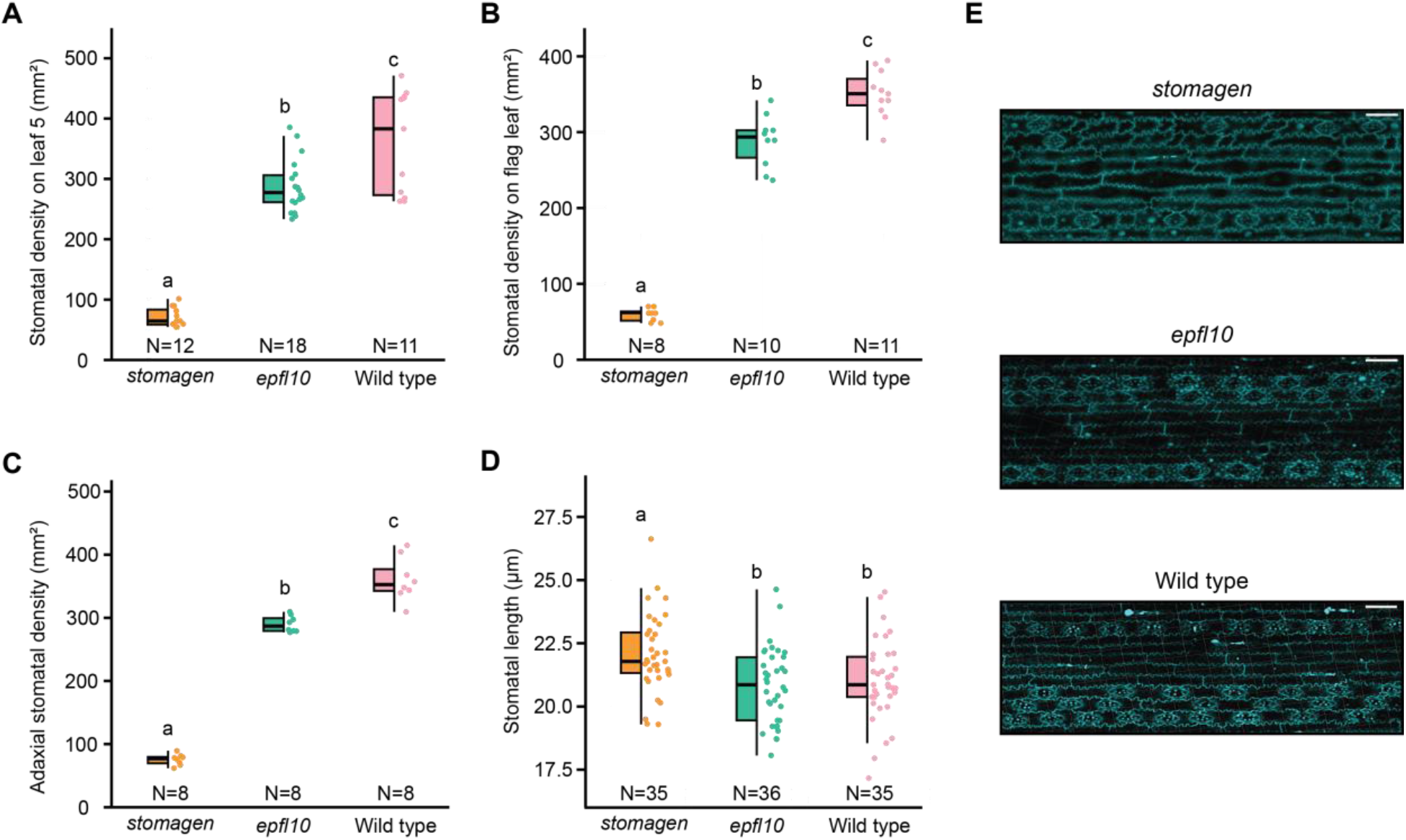
EPFL10 is a weak positive regulator of stomatal development in Nipponbare (*Oryza sativa* spp. Japonica)**. A**-**C**, Stomatal density of *epfl10, stomagen*, and wild type. **A**, Stomatal density of the fifth fully expanded true leaf. Stomatal densities of 21-day old plants were measured. **B**, Stomatal density of the flag leaf on the primary tiller during grain filling. Flag leaves of 55-day old plants were measured. **C**, Adaxial stomatal densities of the fifth fully expanded true leaf on 21-day old plants. **D**, Stomatal length of *epfl10, stomagen*, and wild type. **E**, Representative confocal images of *epfl10, stomagen*, and wild type. Images were taken of 55-day-old leaves stained with propidium iodide. **A**-**D**, Graphs are box-and-whisker plots where the center horizontal indicates the median, upper and lower edges of the box are the upper and lower quartiles and whiskers extend to the maximum and minimum values within 1.5 interquartile ranges. Letters indicate a significant difference between means (P<0.05, one-way ANOVA Duncan post-hoc test).

### Stomatal density reductions are concomitant with gas exchange reductions in *stomagen* but not *epfl10*

The assayed genotypes revealed anatomical level differences in stomatal densities and length. To determine if these developmental differences corresponded to alterations in carbon assimilation (A_n_) and stomatal conductance (g_s_), the efficiency of gas-exchange was measured across genotypes in response to increasing incident light.

At ambient CO_2_ (400 ppm), *epfl10* maintained wild-type levels of A_n_ and g_s_ whereas *stomagen* exhibited reduced gas exchange capacity in light response curves relative to *epfl10* and wild type at all light intensities greater than 1,000 μmol photons m^−2^ s^−1^ (Fig. 3A and 3B). In an independent cohort of plants assayed at saturating light (1,000 μmol photons m^−2^ s^−1^), *stomagen* steady-state A_n_ and g_s_ were lower relative to wild-type and *epfl10* grown in growth chambers (Fig. <u>S</u>). Similar reductions of A_n_ and g_s_ in *stomagen* lines were recapitulated in greenhouse measurements at 1,000 μmol photons m^−2^ s^−1^ (Fig. S1). However, under vegetative drought *stomagen* did not exhibit lowered levels of carbon assimilation at 1,000 μmol photons m^−2^ s^−1^ despite reduced levels of stomatal conductance (Fig. S1c-d).

**Figure 3.**
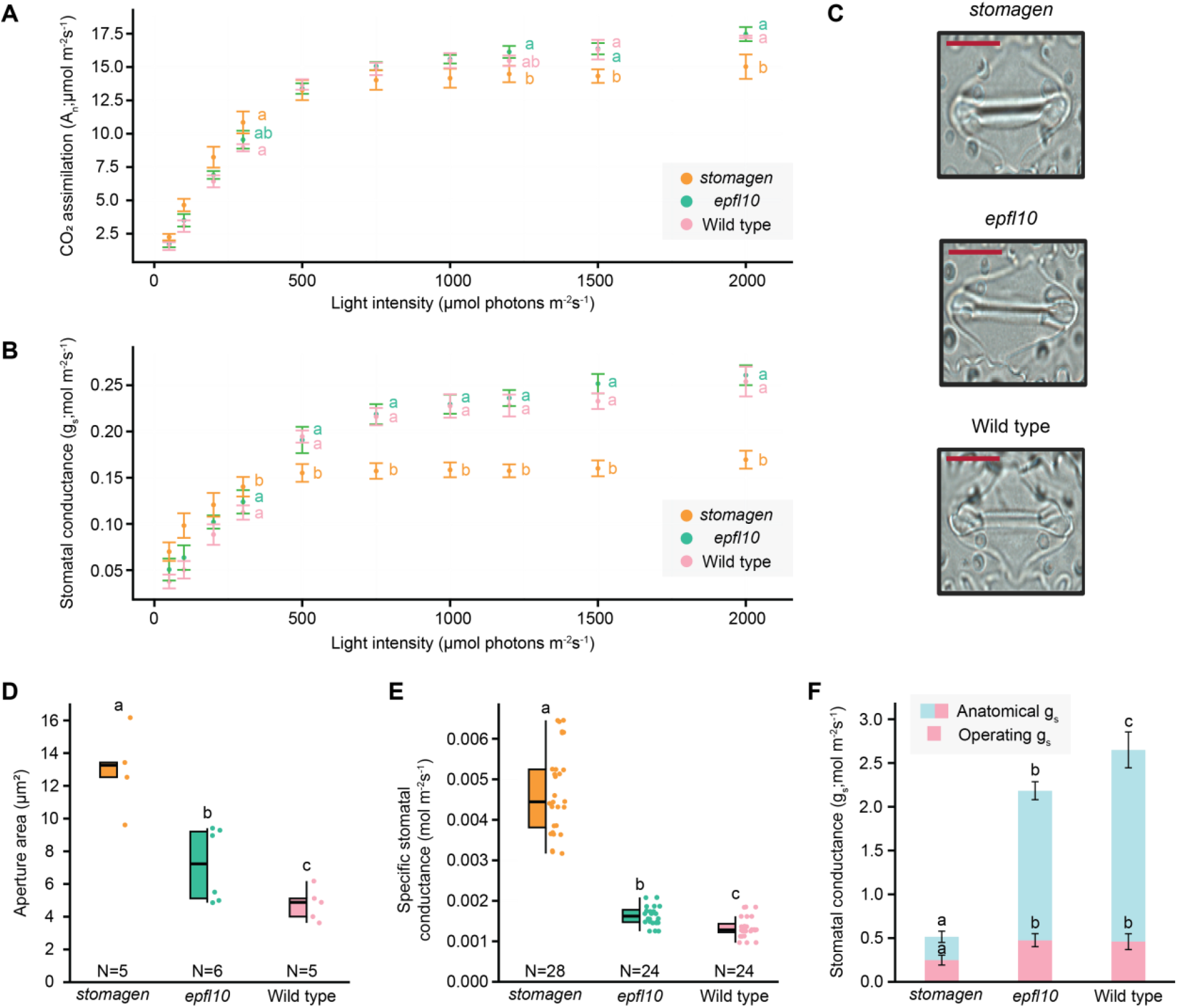
Gas exchange measurements and stomatal pore area measurements in reduced stomatal density backgrounds in Nipponbare (*Oryza sativa* spp. Japonica). **A** and **B**, Carbon assimilation (A) and stomatal conductance (B) measurements of *stomagen, epfl10*, and wild-type across a range of light intensities: 100, 200, 300, 500, 650, 1000, 1200, 1500, and 2000 μmol photons m^−2^ s^−1^ in 32-day-old plants grown in the greenhouse. **C**, Representative images of stomatal pore size variations for *epfl10, stomagen*, and wild type. **D**, Stomatal aperture of *epfl10, stomagen*, and wild type. **E**, Specific stomatal conductance of *epfl10, stomagen*, and wild type. Specific conductance values were calculated by dividing stomatal conductance by the average stomatal density of the probe area of the respective genotype. **F**, Anatomical g_smax_ was calculated for each genotype. Operating g_s_ at 1,500 μmol photons m^−2^ s^−1^ is reported in pink. The percentage of operating to anatomical gs is reported above the pinkbar for each genotype. **B** and **D**,dots represent means and error bars are standard deviations from the mean. **C**, **E** and **F**, in the box- and-whisker plots, the center horizontal indicates the median, upper and lower edges of the box are the upper and lower quartiles and whiskers extend to the maximum and minimum values within 1.5 interquartile ranges. Letters indicate a significant difference between means (P<0.05, one-way ANOVA Duncan post-hoc test).

Measurements of stomatal apertures on the abaxial side of flag leaves indicated that *epfl10* lines maintained a larger stomatal aperture than the wild type, and *stomagen* lines exhibited an even greater aperture (Fig. 3C and 3D). Thus, *epfl10* and *stomagen* maintained even greater levels of stomatal conductance per individual stoma mediated by a larger aperture in well-watered conditions (Fig. 3E). Increased stomatal aperture at reductions in stomatal density of 25% but not 80% were able to sustain wild-type levels of steady-state A_n_ and g_s_.

### Decreasing stomatal density increases ratio of theoretical stomatal conductance to operating stomatal conductance

Despite similar operating stomatal conductance, *epfl10* and WT significantly differ in their theoretical maximum stomatal conductance. This is in contrast to *stomagen* which had both significantly lower operating and anatomical stomatal conductance. Thus, reductions in stomatal density result in a stomatal conductance that operates at a greater proportion of its theoretical maximum for both *stomagen* and *epfl10*, though the latter is able to compensate to sustain WT gas exchange (Fig 2A, 2B).

### *epfl10* maintains wild-type thermoregulation and yield while increasing water conservation

Measurements of A_n_ and g_s_, and g_smax_ revealed differences in the regulation of gas exchange among wild type, *epfl10*, and *stomagen*. To further resolve the lifetime implications of these edits, we assessed differences in thermoregulatory, yield, and water conservation capacities.

Thermal imaging was used to assess evaporative cooling in lines with altered stomata. In well-watered conditions, *stomagen* lines were warmer on average than wild-type, whereas *epfl10* leaf temperatures were wild-type like or cooler (Fig. 4A). No difference in leaf temperature was detected across genotypes during vegetative drought. To test the impact of reduced stomatal densities on water conservation, daily water loss was measured over the course of a week beginning with 70 days after germination. Despite differences in thermoregulation, *stomagen* and *epfl10* both conserved greater volumes of water in a week by 48 mL and 83 mL, respectively (Fig. 4D).

**Figure 4.**
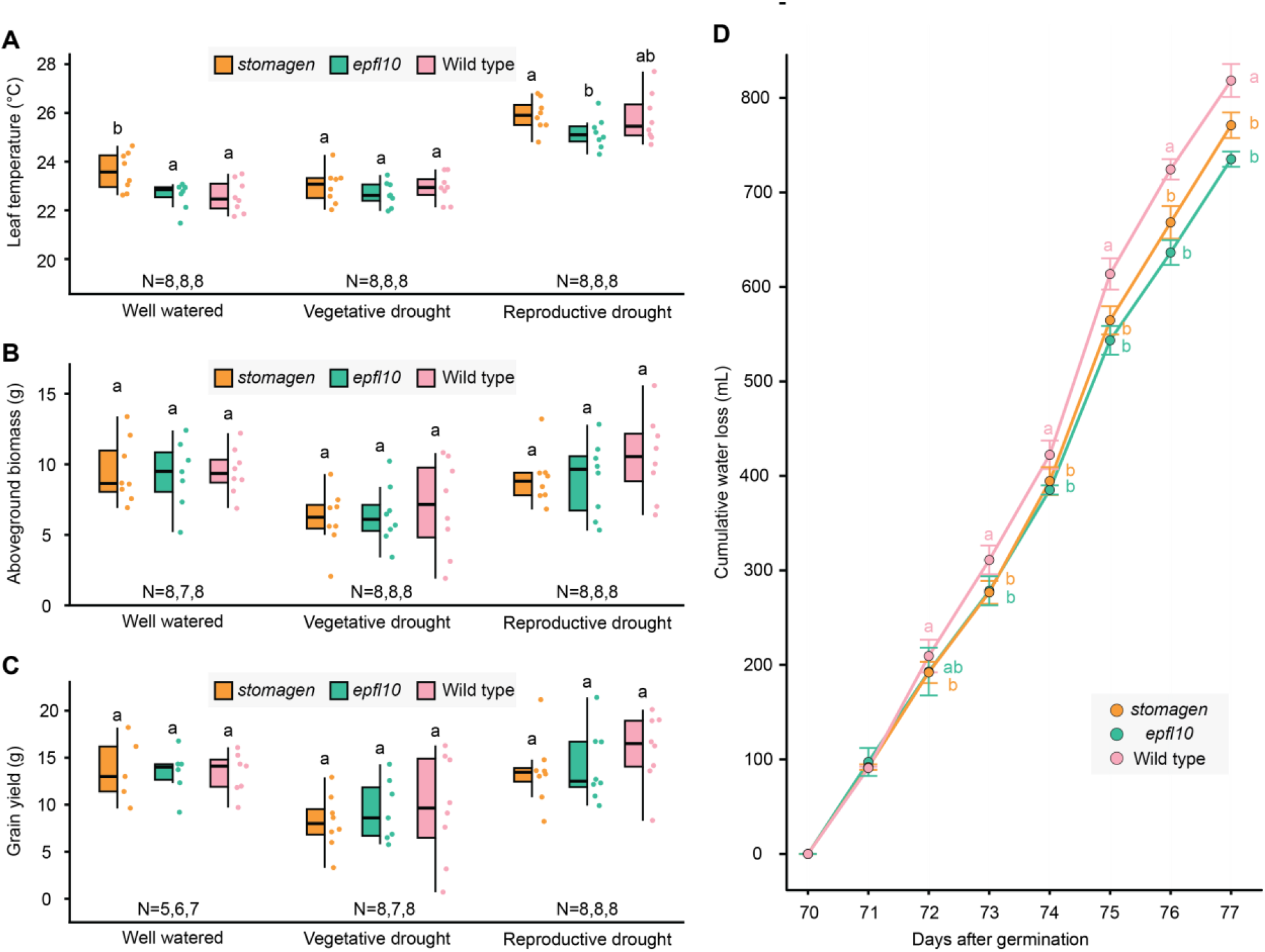
Stomatal density reductions influence thermoregulation and water conservation but not greenhouse yield in Nipponbare (*Oryza sativa* spp. Japonica). **A**, Leaf temperatures of *stomagen, epfl10*, and wild-type in well-watered, vegetative drought, and reproductive drought. **B**, Aboveground biomass of *stomagen, epfl10*, and wild-type in well-watered, vegetative drought, and reproductive drought. **C**, Grain yield of *stomagen, epfl10*, and wild-type in well-watered, vegetative drought, and reproductive drought. **A**-**C**, Graphs are box-and-whisker plots where the center horizontal indicates the median, upper and lower edges of the box are the upper and lower quartiles and whiskers extend to the maximum and minimum values within 1.5 interquartile ranges. **D**, Cumulative water loss of *stomagen, epfl10*, and wild type from days 70-77 after germination. The dots represent means and error bars are standard errors from the mean. Letters indicate a significant difference between means (P<0.05, one-way ANOVA Duncan post-hoc test).

*epfl10* exhibited a range of phenotypes that suggested an increased fitness under drought via moderate stomatal density reductions. This phenotype contrasted with *stomagen*, which may be less optimal for high-yielding production due to large reductions in A_n_ and g_s_. To assess the impacts of stomatal modifications on potential crop performance, yield trials were conducted using three watering regimes in the greenhouse. However, in well-watered, vegetative drought, and 4-day reproductive drought conditions, there was no significant difference in grain yield or aboveground biomass among genotypes (Fig. 4B,C).

### Paralogs of STOMAGEN in other species may also be targets for tuning stomatal density

The potential of paralog editing for tuning stomatal density, as demonstrated in rice *epfl10* lines, can be extended to other crop species. Further phylogenetic investigation of the duplication of STOMAGEN among angiosperms revealed an additional putative family-level STOMAGEN duplication in the Asteraceae (Fig. 5A). Species level duplications were noted in carrot (*Daucus carota*), date palm (*Phoenix dactylifera*), Balbis banana (*Musa balbisiana*), and *Carex littledalei* based on the species included in this study.

**Figure 5.**
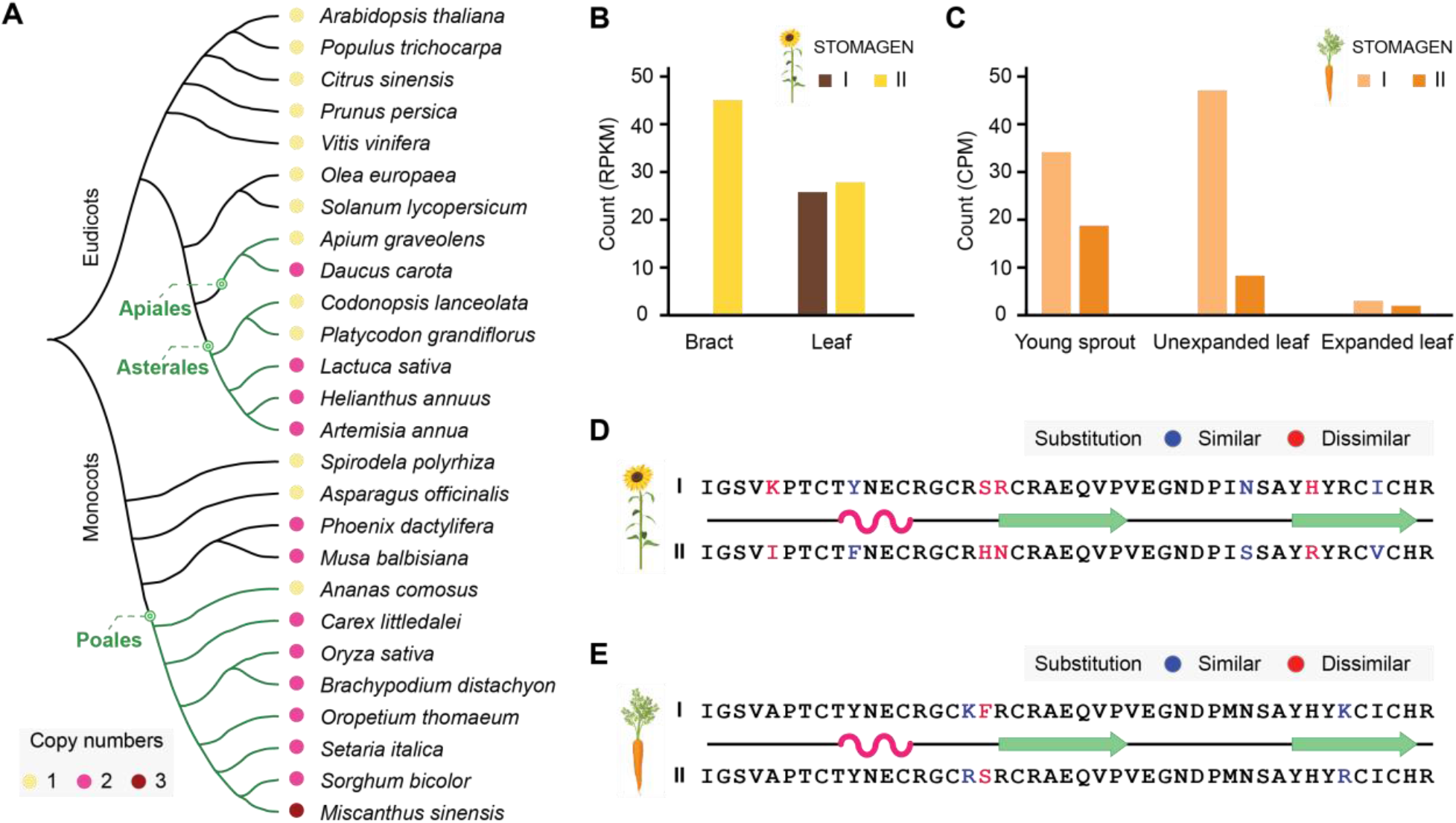
Multiple STOMAGEN duplication events in angiosperms provide new editing targets. **A**, A dendrogram of angiosperm species and copy number variations of STOMAGEN family members. STOMAGEN copy number indicated by colors in legend. **B** and **C**, Expression patterns of the paralogous STOMAGENs in (B) sunflower (*Helianthus anuus*) and (C) carrot (*Daucus carota*) in tissues assayed most relevant to gas exchange. **D** and **E**, The C-terminal active STOMAGEN sequences with non-synonymous substitutions highlighted for the paralogs in sunflower (D) and carrot (E). Substitutions to similar and dissimilar amino acids based on BLOSUM62 are indicated with blue and red The secondary structure annotation originates from the experimentally determined solved structure of STOMAGEN (PDB: 2LIY) ^26^. Alpha helix is represented by pink and beta sheets by green.

Similar to rice STOMAGEN and EPFL10, the paralogs of STOMAGEN in carrot and sunflower are differentially expressed in tissues most relevant to gas exchange suggesting regulatory divergence after the duplication event (Fig 5B, C). Unlike the rice STOMAGEN duplicates, both sunflower and carrot paralogs exhibit less relative sequence variation (Fig. 5D,E). Regardless of functional divergence, duplicated STOMAGEN copies may provide genetic material for optimization of expression in organisms where multiple copies exist.

## Discussion

Drought is the most severe and widespread environmental stressor in South and Southeast Asia ^73^. Application of gene editing in crops for climate change could serve as a potent mechanism for realizing actual technology transfer to growers. Few gene editing applications for addressing abiotic stressors have been reported to date ^74^. Preceding transgenic manipulations of epidermal patterning factors generated rice with improved water conservation and maintenance of yields in greenhouse conditions; however, these lines exhibited reductions in A_n_ and g_s_ ^30,31,33^. Exploration of the rice STOMAGEN paralog, EPFL10, was undertaken as a means of generating novel, transgene-free variation in rice stomatal density. The evolutionary and regulatory features of EPFL10 suggested it as a promising target for engineering drought resilience.

Variation in stomatal density introduced by single-gene knockouts of EPFL10 and STOMAGEN provided a basis for further physiological exploration. Stomatal conductance and photosynthesis of *stomagen* but not *epfl10* were lower at all light intensities greater than 1,000 μmol photons m^−2^ s^−1^ relative to wild-type. In field conditions, photosynthetic flux densities regularly exceed 1,000 μmol photons m^−2^ s^−1^ at midday, coinciding with the hottest parts of the day ^75^. Sustaining photosynthesis may work synergistically with optimizing thermoregulation as *stomagen* exhibits reduced gas exchange simultaneous with warmer temperatures in well-watered conditions whereas *epfl10* maintained wild-type levels of thermoregulations (Fig. 3A, 3B, 4A). In accordance with anticipated global warming caused by climate change, maintained or improved thermoregulation will be vital for crop agronomic performance ^4^.

Larger stomatal apertures were measured in *stomagen* and *epfl10* lines relative to wild type (Fig. 3E). Pore area adjustments in *stomagen* lines were unable to physiologically compensate for large reductions in stomatal densities, unlike *epfl10* lines, which maintained wild-type levels of A_n_ and g_s_ (Fig. 3A-D). The theoretical maximum stomatal conductance of grass stomata greatly exceeds the measured stomatal conductance ^33,43^. *epfl10* maintained high levels of specific stomatal conductance concurrent with overall A_n_ and g_s_ (Fig. 3A-D, 3F, 4D). Enhancing specific stomatal conductance in a reduced density background may thus provide a promising mechanism for maintaining photosynthetic capacities simultaneous with water-use efficiency ^43,76^.

No detectable differences were found in greenhouse grain or biomass yield among the reported genotypes under periods of vegetative or reproductive drought. However, there is a reasonable basis to prioritize maintenance of wild-type gas exchange for maximizing yield. Other literature suggests that high levels of leaf photosynthesis and stomatal conductance are linked to higher yields among C3 crops ^36–41^, and it is possible that more severe drought stress or in-field validation may resolve differences between genotypes.

Independent of biomass and yield, *epfl10* provides a potent demonstration of the capacity to reduce stomatal densities and increase soil water conservation without concomitant reductions in gas exchange essential for optimal crop performance at steady-state conditions. It is noteworthy that *stomagen* lines conserved water to a similar extent as *epfl10* despite having much greater reductions in stomatal density. Larger stomata typically exhibit slower dynamic responses ^77^. Slower rates of environmental response in fluctuating light or environmental conditions may account for a non-linear relationship between stomatal density and water conservation. Our data indicate that fewer stomata with larger apertures offer comparable water conservation properties as lines possessing greater numbers of stomata with smaller apertures. It is still unclear how more severe and field-relevant drought stresses may affect water loss and assimilation when specific stomatal conductance is limiting.

Knockout of the STOMAGEN paralog *epfl10* builds on other work that has shown that null alleles of paralogs can achieve desirable phenotypic outcomes ^49,78^. Rice *EPFL10* had reduced expression levels in relevant tissues and significant peptide level differences from *STOMAGEN*, reinforcing ex-situ evidence that EPFL10 may have evolved a weaker overall function in stomatal development (Fig. 1). A similar exploration of *STOMAGEN* duplicates in other organisms was used to identify other candidate species for paralog editing. STOMAGEN copy II in sunflower and STOMAGEN copy I in carrot also each exhibit lower expression that may substantiate these genes as targets for moderate reductions in stomatal density similar to EPFL10 in rice. Gene editing of paralogs can provide a convenient gene-editing target for quantitative variation in traits like stomatal density.

Peptide variation of the STOMAGEN duplicate in rice (EPFL10) exceeded that of carrot and sunflower (Fig 5). It is enticing to consider a scenario where the significant sequence variation underlying OsEPFL10 at the family and genus level may evidence selection against functional redundancy of STOMAGEN (Fig. 1C). Thus, both decreases in expression (observed in all duplicates) and significant peptide variation (observed in rice EPFL10) may underlie functional natural variation that knocked down stomatal density to restore gene dosage imbalances incurred by stochastic STOMAGEN duplication. Further functional characterization in diverse STOMAGEN duplicates is necessary to resolve their role and costs to plant fitness. Paralog editing of stomatal density and other traits may be a promising avenue to accelerate the production of climate-adapted varieties without the constraints of transgenic germplasm.

To this aim, we report here modest stomatal density reductions in *epfl10* mutants with no associated decreases in stomatal conductance or carbon assimilation. *epfl10* lines maintained wild-type physiological capacities of stomatal conductance, carbon assimilation, thermoregulation, and yield while also conserving more water than wild type. These attributes could contribute to improved climate resilience in current and future conditions where water is limiting, and temperatures are increased. Field-based investigations of *epfl10* and *stomagen* will further resolve the agronomic utility of these edited rice lines.

## Acknowledgements

We would like to acknowledge Denise Schichnes at the Biological Imaging Facility for assistance with confocal microscopy. We also acknowledge the support of Christina Wistrom and all greenhouse staff who provided excellent care of our greenhouse trials. Finally, we would like to acknowledge Julie Gray and Robert Caine who provided helpful feedback on project concept and water loss experimental design.

## Author Contributions

NGK developed project idea and coordinated research efforts. NGK constructed gene-editing vectors with assistance from DD. MT and JT with oversight from MJC produced *epfl10* and *stomagen* lines, respectively. KS generated dendrogram, gene tree, all peptide models, and designed all figures. KS with assistance from NGK determined expression levels in carrot and sunflower. NGK phenotyped stomatal densities, stomatal aperture, and stomatal size with assistance from AGC, SL, and LL. GAG repeated stomatal density phenotyping independently to confirm results. NGK phenotyped water loss and yield. NGK and DPT conducted gs and A assays. DPT captured thermal images. NGK drafted the manuscript with edits from DPT, KS and KKN. KKN provided LICOR equipment, thermal imaging equipment, and technical expertise. BJS provided feedback on experimental design and the manuscript as well as the facilities to carry out experiments.

