## Supporting Figures 1-6 for "Paralog editing tunes rice stomatal density to maintain photosynthesis and improve drought tolerance"

**Figure S1:**

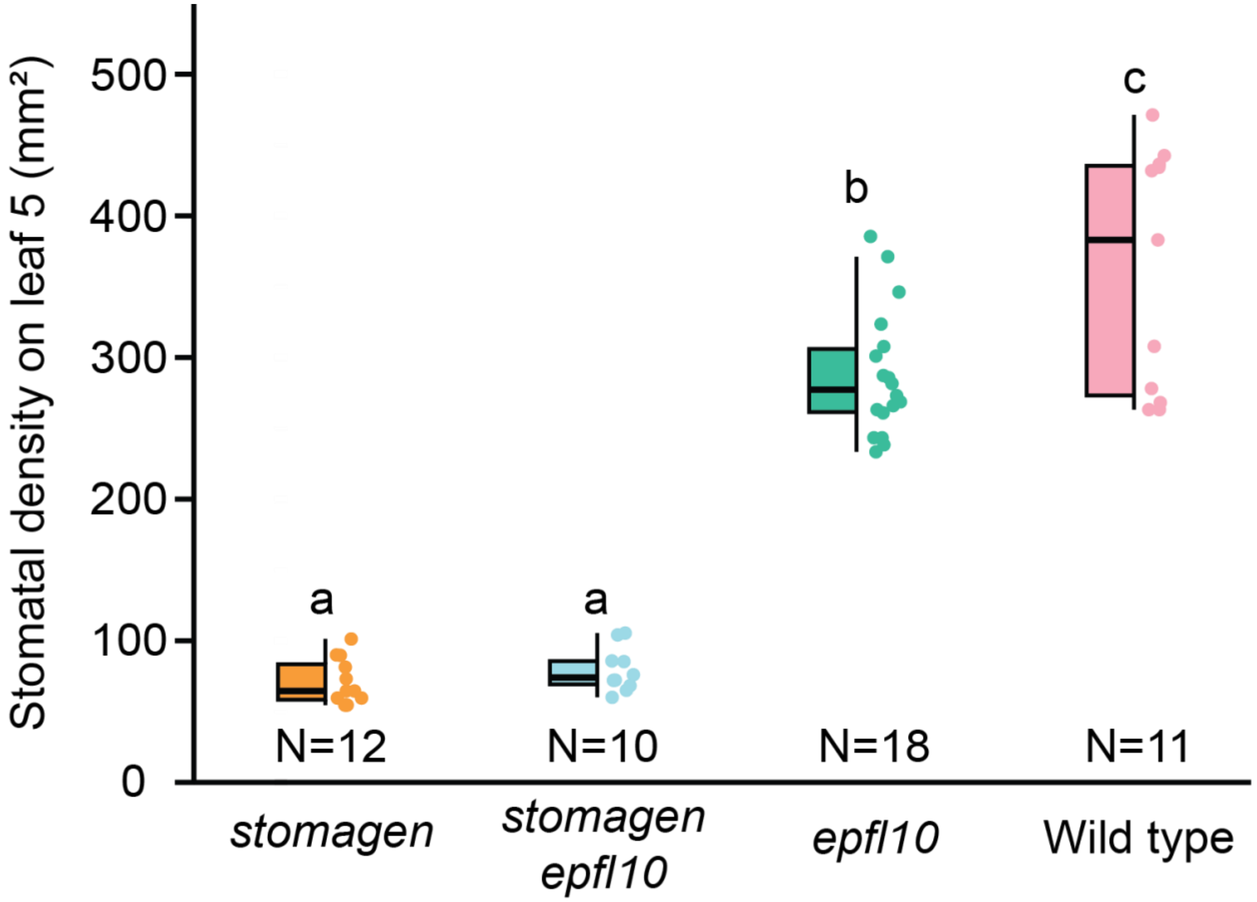


**Figure S2**

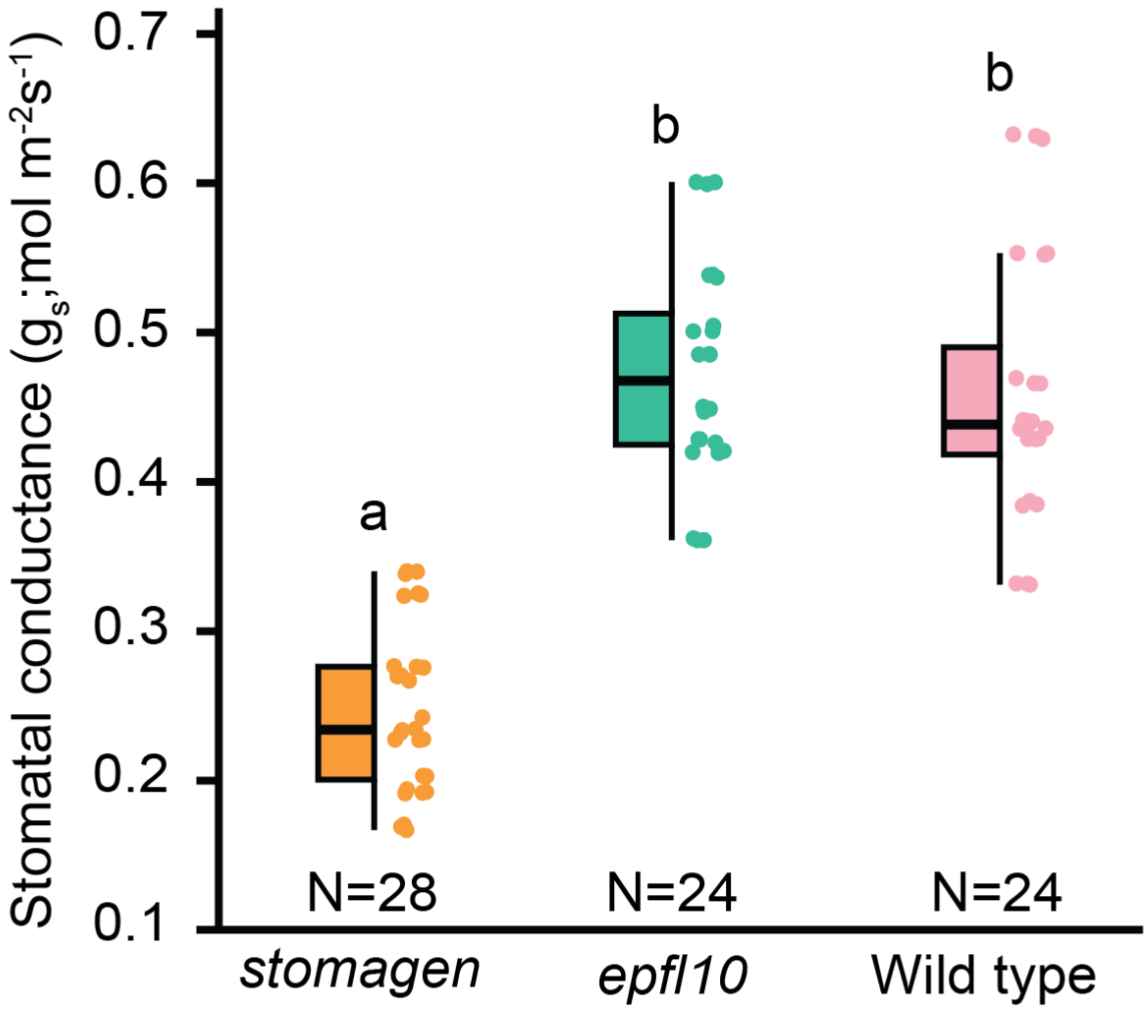
Stomatal conductance (gs;mol m-2s-1)
0.7
0.6
0.5
0.4
0.3
0.2
0.1
a
b
b
N=28
N=24
N=24
stomagen
epfl10
Wild type

**Figure S3**

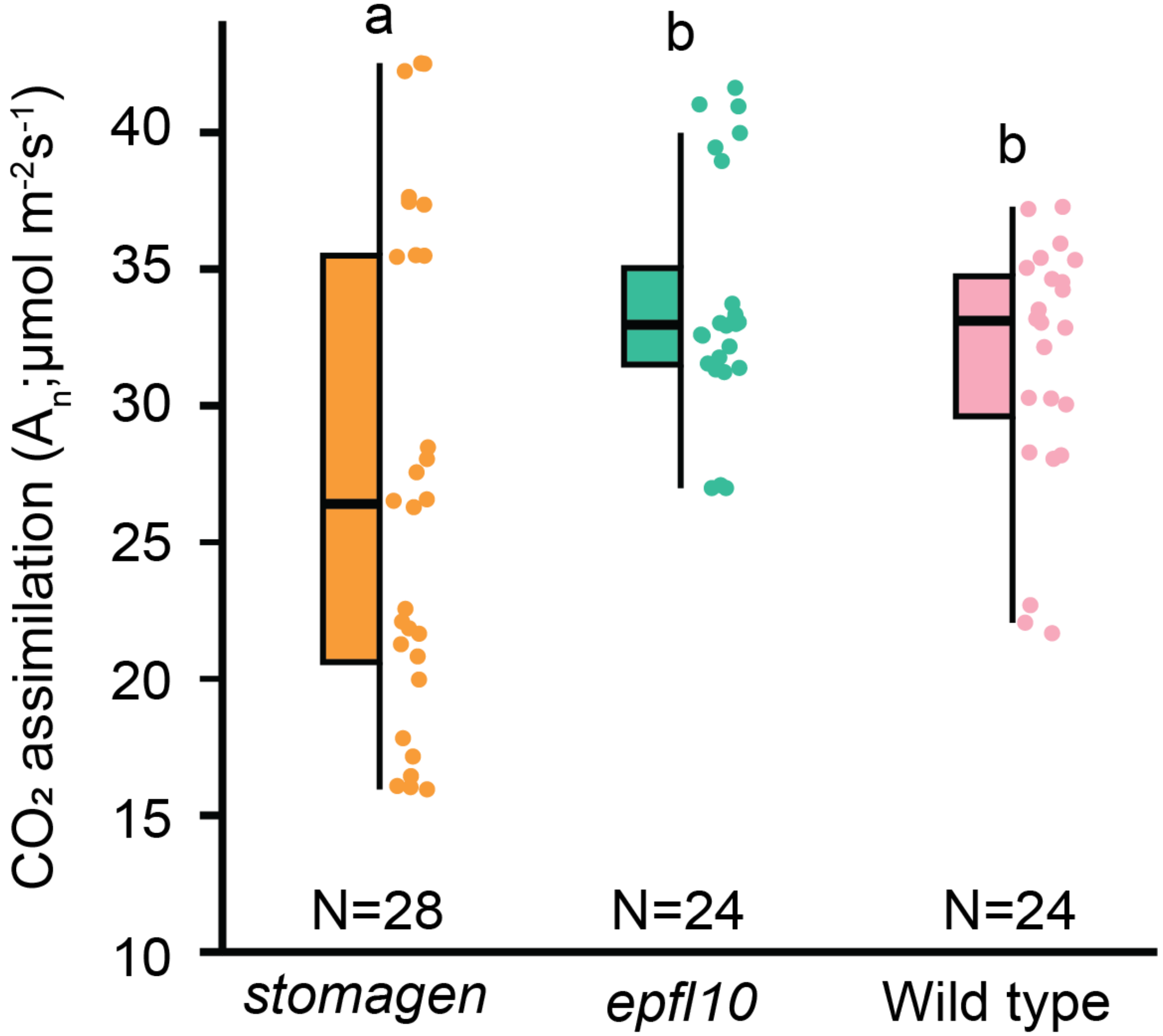
a
b
b
CO₂ assimilation (A_n;µmol m-2s-1)
40
35
30
25
20
15
10
N=28
N=24
N=24
stomagen
epfl10
Wild type

**Figure S4:**

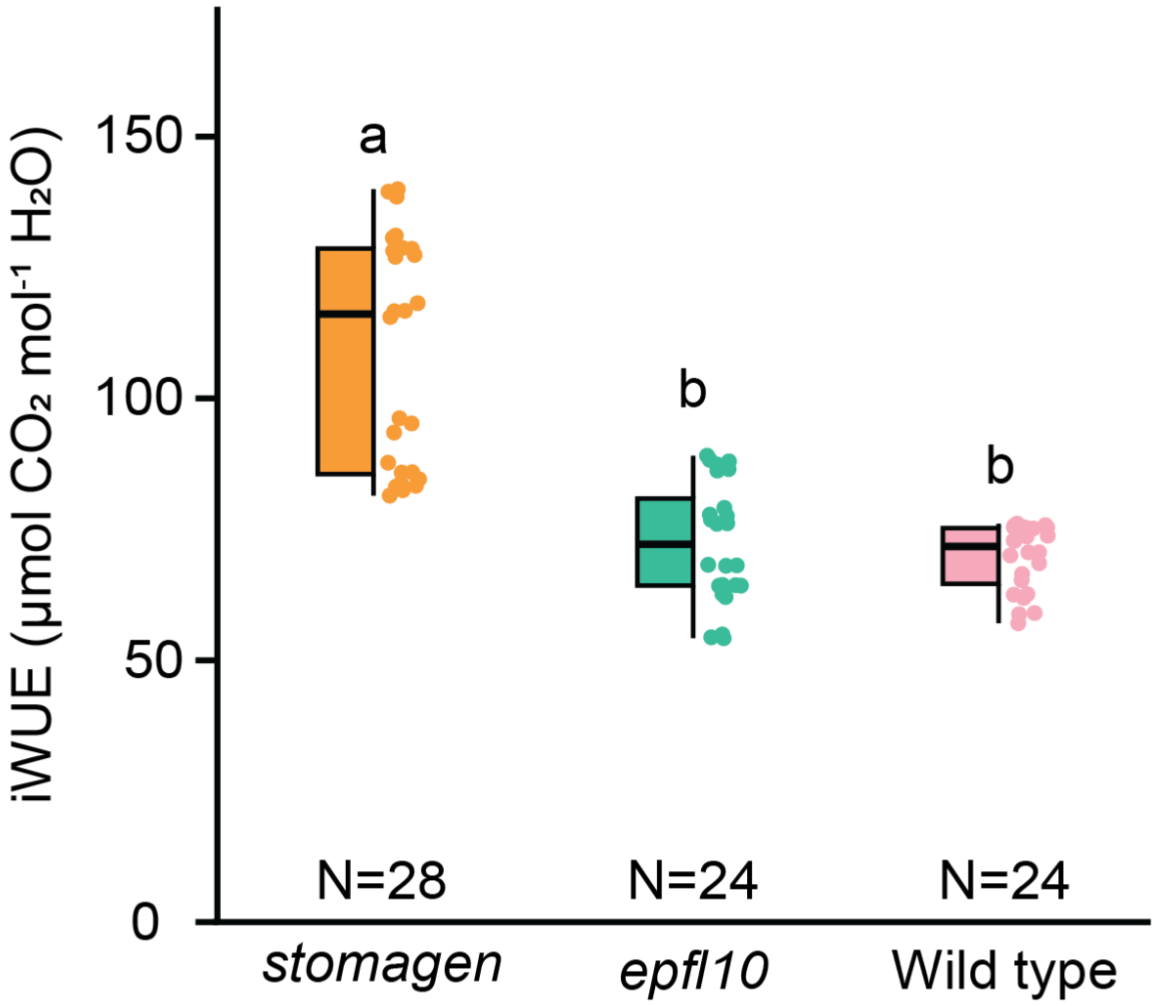
iWUE (µmol CO₂ mol⁻¹ H₂O)
150
100
50
0
a
b
b
N=28
N=24
N=24
stomagen
epfl10
Wild type

**Figure S5:**

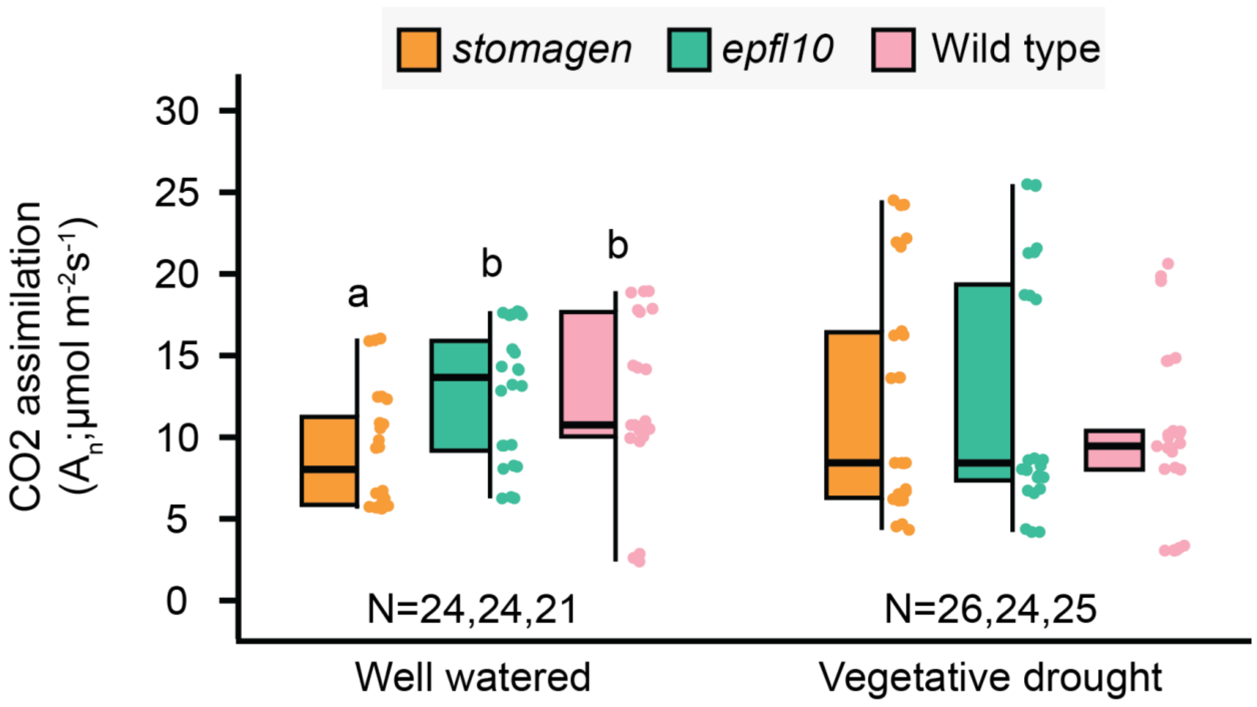
stomagen
epfl10
Wild type
CO2 assimilation
(A_n;µmol m^-2s^-1)
30
25
20
15
10
5
0
a
b
b
N=24,24,21
N=26,24,25
Well watered
Vegetative drought

**Figure S6:**

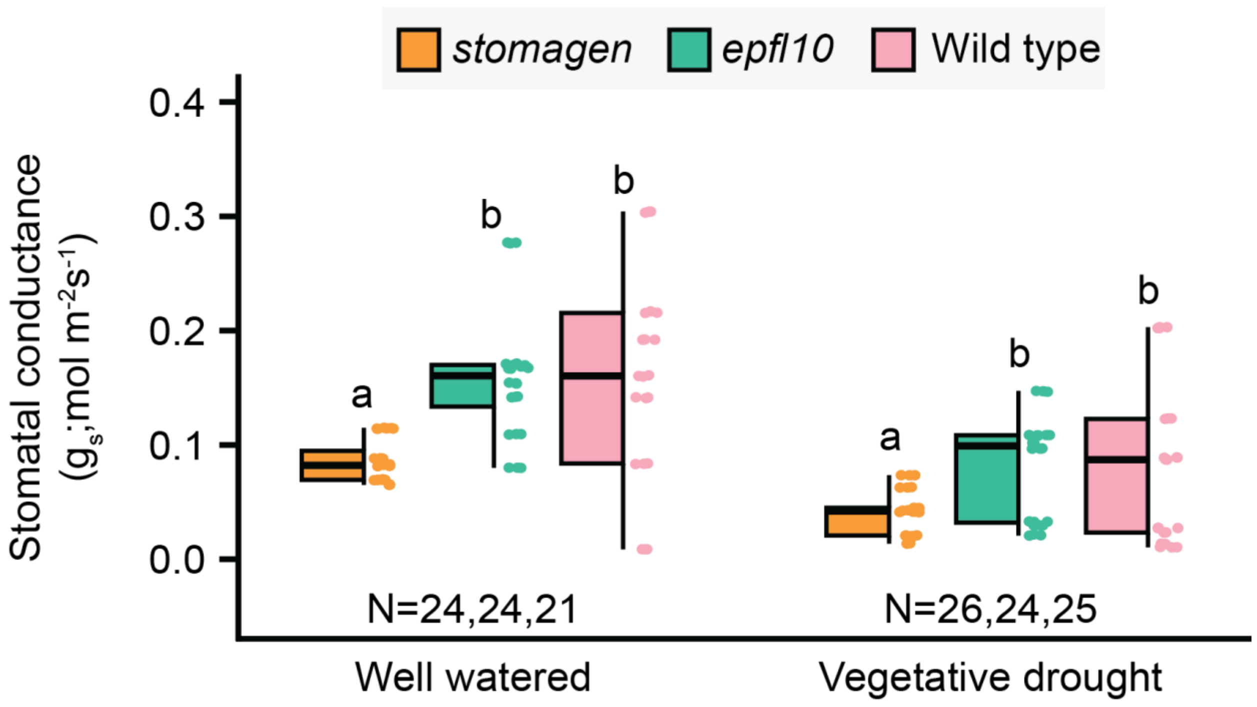

stomagen
epfl10
Wild type
Stomatal conductance
($g_s$; mol $m^{-2}s^{-1}$)
0.4
0.3
0.2
0.1
0.0
a
b
b
a
b
b
N=24,24,21
N=26,24,25
Well watered
Vegetative drought

**Figure S1.)** Stomatal density of the fifth fully expanded true leaf. Stomatal density measurements of *stomagen, epfl10; stomagen; epfl10;* and WT taken on 21-day old plants grown in the growth chamber. In this box-and-whisker plot the center horizontal indicates the median, upper and lower edges of the box are the upper and lower quartiles and whiskers extend to the maximum and minimum values within 1.5 interquartile ranges. Letters indicate a significant difference between means ($P<0.05$, one-way ANOVA Duncan post-hoc test). Measurements were taken on full expanded leaf 5 of 28-day-old plants grown in well-watered conditions plants using an infrared gas analyzer (LI6800XT, LI-COR, Lincoln, NE, USA) with chamber conditions set to: light intensity 1000 µmol photons $m^{-2}$ $s^{-1}$ (90% red light, 10% blue light); leaf temperature 27°C; flow rate 500 µmol $s^{-1}$; relative humidity 40%; and $CO_2$ concentration of sample 400 µmol $mol^{-1}$.

and vegetative drought All data collected were adjusted according to leaf area within the gas exchange chamber. Intrinsic water use efficiency (iWUE) in Figure S3 was calculated by dividing photosynthesis by stomatal conductance for each biological replicate. Specific stomatal conductance in Fig 3f was calculated by dividing stomatal conductance by the average number of stomata within the probe area.

Physiological assays in Figures 3A, 3B were conducted on fully expanded leaf 5 of 28-day old plants using an infrared gas analyzer (LI6800XT, LI-COR, Lincoln, NE, USA) with chamber conditions set to: light intensity 1000 µmol photons $m^{-2}$ $s^{-1}$ (90% red light, 10% blue light); leaf temperature 27°C; flow rate 500 µmol $s^{-1}$; relative humidity 40%; and $CO_2$ concentration of sample 400 µmol $mol^{-1}$.

**Figure S2.)** Carbon assimilation measurements of *stomagen*, *epfl10*, and wild-type at 1000 µmol photons $m^{-2}$ $s^{-1}$ grown in the growth chamber. In this box-and-whisker plot the center horizontal indicates the median, upper and lower edges of the box are the upper and lower quartiles and whiskers extend to the maximum and minimum values within 1.5 interquartile ranges. Letters indicate a significant difference between means ($P<0.05$, one-way ANOVA Duncan post-hoc test). Measurements were taken on full expanded leaf 5 of 28-day-old plants grown in well-watered conditions plants using an infrared gas analyzer (LI6800XT, LI-COR, Lincoln, NE, USA) with chamber conditions set to: light intensity 1000 µmol photons $m^{-2}$ $s^{-1}$ (90% red light, 10% blue light); leaf temperature 27°C; flow rate 500 µmol $s^{-1}$; relative humidity 40%; and $CO_2$ concentration of sample 400 µmol $mol^{-1}$.

**Figure S3.)** Stomatal conductance measurements of *stomagen*, *epfl10*, and wild-type at 1000 µmol photons $m^{-2}$ $s^{-1}$ grown in the growth chamber. In this box-and-whisker plots the center horizontal indicates the median, upper and lower edges of the box are the upper and lower quartiles and whiskers extend to the maximum and minimum values within 1.5 interquartile ranges. Letters indicate a significant difference between means ($P<0.05$, one-way ANOVA Duncan post-hoc test)

**Figure S4**. Instantaneous water-use efficiency of *epfl10* and *stomagen* lines in Nipponbare (*Oryza sativa* spp. Japonica) grown in the growth chamber. All graphs in figure are box-and-whisker plots where the center horizontal indicates the median, upper and lower edges of the box are the upper and lower quartiles and whiskers extend to the maximum and minimum values within 1.5 interquartile ranges. Letters indicate a significant difference between means ($P<0.05$, one-way ANOVA Duncan post-hoc test). Measurements were taken on full expanded leaf 5 of 28-day-old plants grown in well-watered conditions. Intrinsic water use efficiency (iWUE) in Figure S3 was calculated by dividing photosynthesis by stomatal conductance for each biological replicate.

**Figure S5**. Measurements of carbon assimilation *epfl10* and *stomagen* lines in Nipponbare (*Oryza sativa* spp. Japonica) grown in the greenhouse under two watering regimes, well-watered, and vegetative drought. In this box-and-whisker plots the center horizontal indicates the median, upper and lower edges of the box are the upper and lower quartiles and whiskers extend to the maximum and minimum values within 1.5 interquartile ranges. Letters indicate a significant difference between means ($P<0.05$, one-way ANOVA Duncan post-hoc test). Measurements were taken on full expanded leaf 5 of 28-day-old plants grown in well-watered conditions and vegetative drought.

**Figure S6**. Measurements of stomatal conductance *epfl10* and *stomagen* lines in Nipponbare (*Oryza sativa* spp. Japonica) grown in the greenhouse under two watering regimes, well-watered, and vegetative drought. In this box-and-whisker plots the center horizontal indicates the median, upper and lower edges of the box are the upper and lower quartiles and whiskers extend to the maximum and minimum values within 1.5 interquartile ranges. Outliers are represented by black dots. Letters indicate a significant difference between means ($P<0.05$, one-way ANOVA Duncan post-hoc test).
